# Pharmacokinetics and efficacy of tank-water administered BRAF-inhibitor dabrafenib in a zebrafish model of BRAF-mutant melanoma

**DOI:** 10.1101/2025.10.12.681850

**Authors:** Jenna Villman, Nikol Dibus, Jonas Huhn, Anna-Mari Haapanen-Saaristo, Ilia Evstafv, Alex Dickens, Oliver Scherf-Clavel, Ilkka Paatero, Kari J. Kurppa

## Abstract

Zebrafish models are widely used to study the biology of BRAF-mutant melanoma. However, long-term treatment of adult fish with small molecule BRAF inhibitors is challenging, limiting the usefulness of this model to study treatment-induced effects in melanoma biology. In addition, pharmacokinetic studies on small molecule inhibitors in zebrafish that could inform rational dosing strategies, are largely lacking. Here, we have assessed the pharmacokinetics, metabolism and efficacy of continuous tank water -administered BRAF-inhibitor dabrafenib in adult zebrafish. Our results demonstrate that dabrafenib is quickly absorbed from the tank water, reaching efficacious plasma levels within one hour following treatment, but also shows fast elimination kinetics with a half-life of 1.6 hours. We could detect most of the human metabolites of dabrafenib in zebrafish, suggesting that dabrafenib metabolism in zebrafish follows a similar process as in humans. Continuous tank water -administered dabrafenib led to therapeutically relevant steady-state plasma levels that inhibited the BRAF-driven signaling and growth in zebrafish melanoma cells *in vitro*, and resulted in robust *in vivo* efficacy in a genetic zebrafish model of BRAF-mutant melanoma, with no apparent toxicity. Together, our results demonstrate that continuous tank water -administered dabrafenib provides a feasible, efficient, and well-tolerated dosing strategy to study treatment-related effects in zebrafish models of BRAF-mutant melanoma. We expect that tank water-administration may also facilitate the dosing of other small molecule inhibitors, especially those with short *in vivo* half-life in zebrafish.

**Highlights:**

- Pharmacokinetic analysis demonstrates fast absorption kinetics and short plasma half-life for tank water -administered dabrafenib in zebrafish
- Dabrafenib is metabolized in zebrafish following a similar metabolic process as in humans
- Tank water -administered dabrafenib provides a feasible, efficient, and well-tolerated dosing strategy to study treatment-related effects in zebrafish models of BRAF-mutant melanoma
- Tank water-administration may facilitate dosing of small molecule inhibitors with short *in vivo* half-life in zebrafish

## 1. Introduction

Melanoma incidence and deaths are constantly rising [1,2]. While local melanoma can be surgically removed and has favorable prognosis, invasive or metastasized melanoma is typically highly aggressive, and the prognosis is poor [2]. In roughly 50% of melanoma cases, the disease is driven by oncogenic mutations in the *BRAF* gene, most often by the BRAF V600E mutation [1]. Melanoma patients with BRAF mutations are clinically treated with BRAF-targeted therapy, such as dabrafenib or vemurafenib, often in combination with MEK inhibitors [1]. While the response rates to BRAF-targeted therapy are typically very good, acquired resistance develops in almost all cases, preventing long-term benefit from therapy.

Zebrafish models have become important tools to understand melanoma genetics and cellular biology, especially in the context of BRAF-mutant melanoma. Since the introduction of zebrafish models of BRAF-mutant melanoma by Patton and others [3] numerous studies have used this model system to provide insight into melanoma development and progression. However, the long-term treatment of adult fish using BRAF-targeted therapy is challenging, limiting the usefulness of this model for studies on adaptive or intrinsic resistance mechanisms to BRAF-targeted therapy.

Previous studies have described the administration of BRAF-inhibitor vemurafenib to adult fish by oral gavage [4], or by pellets of vemurafenib-containing food [5]. However, gavage to a fish of few centimeters in length is challenging and required expertise and may cause severe morbidity in the treated fish if done wrong. Also, continuous and even uptake of drug *via* food is difficult to control, potentially leading to uneven dosing. Furthermore, the understanding of the pharmacokinetics of many small molecule inhibitors, including ones targeting BRAF, in zebrafish is still limited and precludes rational dosing design.

Pharmacological and toxicological studies on larval Zebrafish routinely administer drugs by supplementing the tank water with the study compounds. However, this route of administration is used to dose adult fish in e.g. acute toxicity studies [6] but less commonly used for longer efficacy studies. To date, it has not been used to dose small molecule inhibitors in zebrafish melanoma models. Here, we studied the feasibility, pharmacokinetics, and efficacy of continuous tank water - administered BRAF-inhibitor dabrafenib in adult fish, using a well-established zebrafish model of BRAF-mutant melanoma. Our results demonstrate that dabrafenib administered in the tank water is quickly absorbed into adult fish, reaching efficacious levels within 1 hour of exposure to drug. Furthermore, we demonstrate that dabrafenib is metabolized in zebrafish to same metabolites as in humans and show fast elimination kinetics. Treatment of tumor-bearing fish resulted in rapid and sustained tumor regression, both in melanomas of the skin as well as in ocular melanomas. Together, our results demonstrate that administering dabrafenib directly in tank water provides a feasible, efficient, and well-tolerated method to study treatment-related effects in zebrafish models of BRAF-mutant melanoma.

## 2. Results

### 2.1. Tank water -administered dabrafenib has rapid pharmacokinetics in adult zebrafish and leads to sustained therapeutically relevant plasma levels over long-term treatment

To develop a cost-effective, fast and efficient method to dose adult zebrafish with BRAF-inhibitor dabrafenib over long treatment durations, we assessed the feasibility of directly administering the drug to the fish tank water. To analyze the pharmacokinetics of tank-water -administered dabrafenib in zebrafish, we exposed adult zebrafish to 10 µM dabrafenib in tank water and collected post-mortem blood samples from fish exposed to dabrafenib for different time (Fig. 1A). These blood samples were then subjected to mass spectrometry (MS) analysis for dabrafenib and its active metabolite hydroxy-dabrafenib. Blood sampling was carried out after euthanasia and yielded 0.6 – 6.4 microlitres of plasma per fish, with good correlation to fish weight (Fig. 1B). Despite low volumes, all samples were sufficient for MS analysis (Supplementary Table 1). Based on the analysis, dabrafenib was absorbed rapidly and detected in zebrafish plasma already after 1 hour of exposure (Fig. 1C). Dabrafenib plasma levels peaked at 4 hours post-exposure, after which the levels started to decline, reaching 2.7 +/-2.3 µM at 24 hours post exposure, suggesting active metabolism of dabrafenib in zebrafish. Indeed, we were able to detect dabrafenib metabolite hydroxy-dabrafenib soon after administration of dabrafenib, with similar kinetics to dabrafenib (Fig. 1D). To analyze the elimination of dabrafenib, we transferred zebrafish to fresh water after 24-hour dabrafenib exposure and followed dabrafenib levels in fish plasma over time (Fig. 1A). Dabrafenib was cleared rapidly from zebrafish plasma with a half-life of 1.0 hour (Fig. 1E) and similar fast elimination kinetics (t1/2 1.6h) were observed for hydroxy-dabrafenib (Fig. 1F). These results demonstrate fast absorption and elimination kinetics, as well as active metabolism of tank-water -administered dabrafenib in zebrafish, with therapeutically relevant [7] steady-state plasma levels using 10 µM tank-water concentrations of dabrafenib. Indeed, after 2-week treatment, the dabrafenib concentration in fish plasma was 5.9 +/- 3.3 µM and hydroxy-dabrafenib 0.14 +/- 0.12 µM (Supplementary Table 1), indicating sustained therapeutically relevant levels also over long-term treatment period.

**Figure 1.**
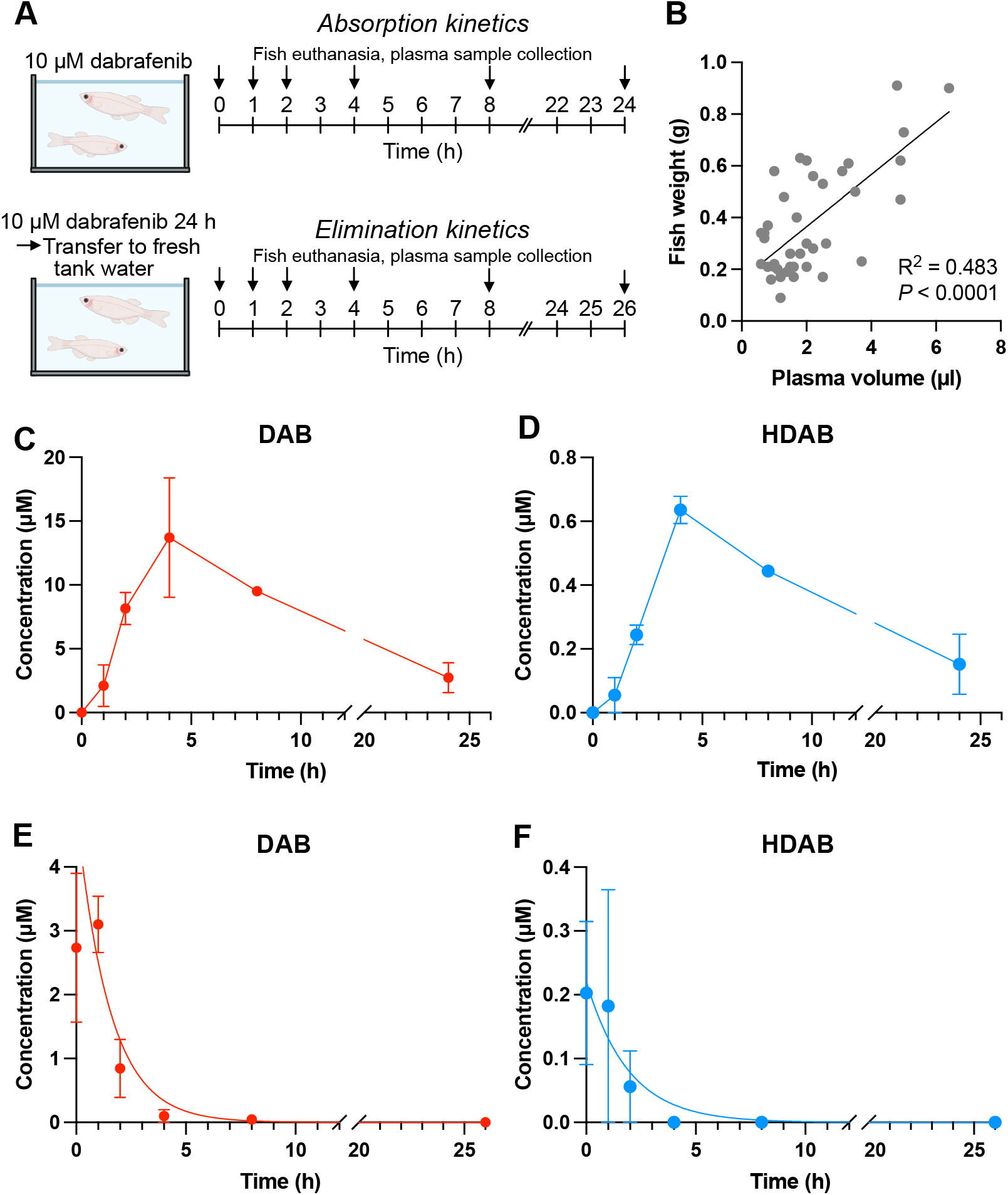
Dabrafenib administered to tank water shows rapid absorption and elimination kinetics in zebrafish. **A)** Schematic overview of sample collection. To monitor absorption, the zebrafish were kept in dabrafenib supplemented water for different times, culled and blood samples collected. To monitor elimination, after 24-h drug exposure, the zebrafish were transferred to water without dabrafenib and culled at different time points. **B)** Terminally collected blood sample volumes correlated with fish weight. Pearson correlation. n= 39 **C** and **D)** Absorption of dabrafenib (DAB) and emergence of hydroxy-dabrafenib (HDAB) were measured from blood samples using mass-spectrometry. Time (h) is time in hours after fish transferred into dabrafenib containing water. **E and F)** Elimination of dabrafenib (DAB) and hydroxy-dabrafenib (HDAB) were measured from blood samples using mass-spectrometry. Time (h) is time in hours after dabrafenib treated fish was transferred into water without dabrafenib.

### 2.2. Dabrafenib is metabolized in zebrafish through a similar metabolic process as in humans

To further analyze the metabolism and distribution of dabrafenib in zebrafish tissues, zebrafish exposed to 10 µM dabrafenib for 24 hours were euthanized, snap-frozen and cryosectioned. Cryosections were analyzed using mass spectrometry imaging (Fig. 2A) for dabrafenib and its main metabolites in humans, hydroxy-dabrafenib, carboxy-dabrafenib, and desmethyl-dabrafenib. In humans, dabrafenib is metabolized by Cytochrome P450 (CYP) enzymes to hydroxy-dabrafenib, and further to carboxy-dabrafenib. Carboxy-dabrafenib can be excreted in bile and urine, or can undergo a non-enzymatic decarboxylation in the gut to desmethyl-dabrafenib [7]. We were able to detect dabrafenib, hydroxy-dabrafenib and desmethyl-dabrafenib in zebrafish tissues but not carboxy-dabrafenib (Fig. 2B), suggesting that dabrafenib undergoes similar metabolism in zebrafish as in humans. While we cannot exclude the possibility of technical limitations in detecting carboxy-dabrafenib, it is possible that carboxy-dabrafenib undergoes fast non-enzymatic conversion to desmethyl-dabrafenib in zebrafish gut. Indeed, the localization of mass spectrometry imaging signal was indicative of distribution of dabrafenib and its detected metabolites especially in the gut, (Fig. 2C-E), although sample tissue sample showed suboptimal integrity. Taken together, these observations suggest that dabrafenib is metabolized in zebrafish through a very similar metabolic process as in humans.

**Figure 2.**
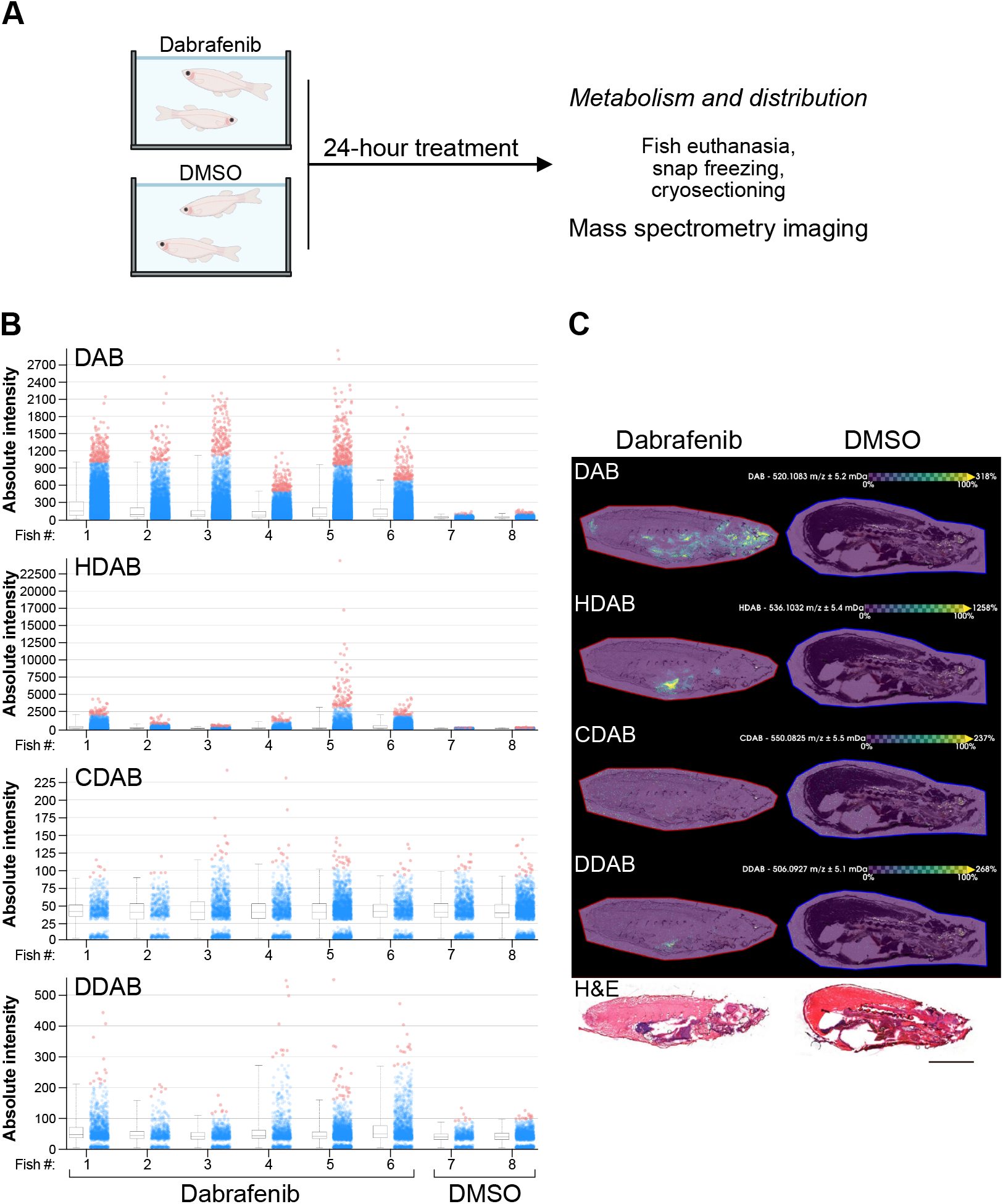
Dabrafenib administered to tank water shows relevant metabolism and distribution in zebrafish. **A)** Schematic overview of sample collection. To monitor distribution, the zebrafish were kept in dabrafenib supplemented water for 24 hours, euthanized and snap-frozen. Frozen samples were cryosectioned sagitally and processed for mass-spectrometry imaging. **B)** Total intensity of dabrafenib (DAB), hydroxy-dabrafenib (HDAB), carboxy-dabrafenib (CDAB) and desmethyl-dabrafenib (DDAB) in cryosections. **C)** Distribution of dabrafenib and its metabolites in zebrafish cryosections. DAB was widely distributed, whereas HDAB and CDAB were localized, presumably in the gut. CDAB levels were very low/absent and it could not be measured with sufficient confidence.

### 2.3. Steady-state dabrafenib plasma levels are sufficient to suppress MAPK signaling and proliferation of BRAF-driven zebrafish melanoma in vitro

Oncogenic mutations in BRAF, such as the BRAF V600E, lead to hyperactivation of the downstream mitogen-activated protein kinase (MAPK) pathway, which fuels the growth of melanoma cells. In order to assess whether the reached steady-state dabrafenib plasma concentrations were sufficient to suppress MAPK signaling and proliferation of BRAF V600E -driven zebrafish melanoma, we analyzed ZMEL1 cells [8] *in vitro*. A western blot analysis of phospho-Erk levels indicated that dabrafenib effectively inhibits MAPK signaling in ZMEL1 cells with IC50 of 75 nM (CI 95% 40-146 nM) (Fig. 3A, B). Consistently, we observed a dose-dependent reduction in ZMEL1 cell growth with an IC50 of 74 nM (95% CI 57-97 nM) (Fig. 3C). Both IC50 values are well below the reached steady state dabrafenib plasma levels of 2.7 µM. Indeed, this concentration led to more than 80% reduction in both phospho-Erk levels and cell proliferation in ZMEL1 cells (Fig. 3B, C). Taken together, the tank-water -administered dabrafenib results in steady-state dabrafenib plasma levels that are able to suppress MAPK signaling and proliferation of BRAF V600E -driven zebrafish melanoma *in vitro*.

**Figure 3.**
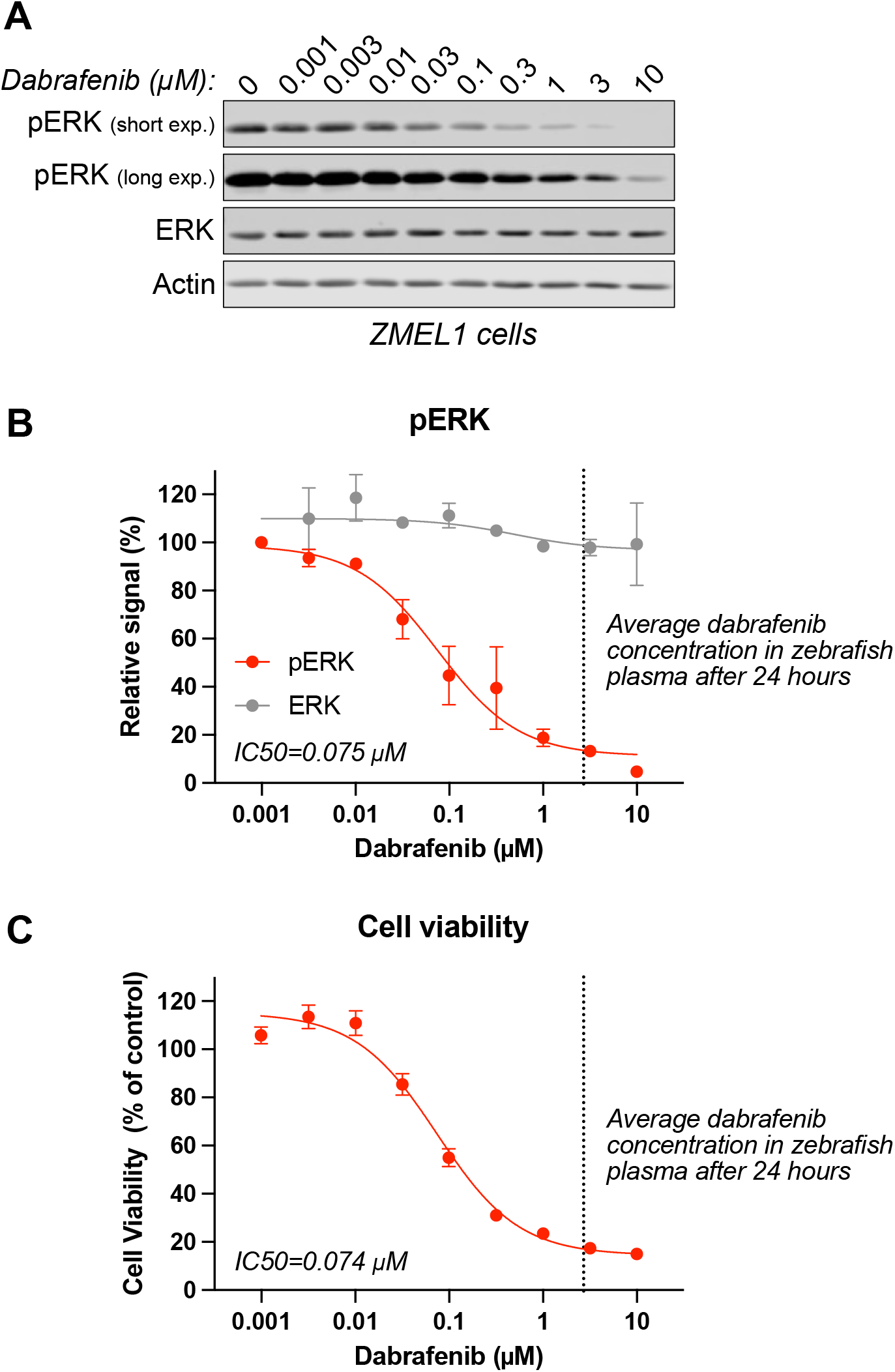
Steady-state dabrafenib plasma levels are sufficient to suppress MAPK signaling and proliferation of BRAF-driven zebrafish melanoma in vitro. **A)** Dabrafenib-induced reduction in MAPK activity in BRAF-driven zebrafish melanoma cells. ZMEL1 cells were treated for 24 hours with the indicated concentrations of dabrafenib, and protein lysates were analyzed for phospho-ERK using western blotting. **B)** Quantification of the results in (A). The data represent the average of two independent experiments and shown as mean ± SD **C)** Dabrafenib-induced reduction in ZMEL1 cell proliferation. The cells were treated with the same dabrafenib concentrations as in (A) for 72 hours, followed by viability measurement. The data shown represent the average of three independent experiments and is shown as mean ± SEM. The steady-state dabrafenib plasma concentration shown in Fig 1C is indicated as a vertical dashed line in (B) and (C).

### 2.4. Tank water -administered dabrafenib results in rapid and sustained tumor regression in zebrafish model of BRAF-mutant melanoma

To assess the *in vivo* efficacy of tank water -administered dabrafenib in a genetic zebrafish model of BRAF-mutant melanoma, we treated melanoma tumor -bearing zebrafish for two weeks with 10 µM dabrafenib. Dabrafenib was administered into tank water and the fish were weighed, imaged and the dosing replenished every 3-4 days (Fig. 4A). Consistent with our *in vitro* analyses, the tank water - administered dabrafenib demonstrated robust efficacy *in vivo*, with 9 of 13 (69%) skin melanoma tumors responding to treatment (Fig. 4B), which is in line with clinical studies with BRAF-inhibitors in human patients with BRAF-mutant melanoma [9,10] All of the nine responding tumors regressed more than 50% in size, with near complete responses in three cases. Overall, tank water -administered dabrafenib resulted in significant reduction in skin melanoma tumor burden in the treated fish, compared to DMSO-treated controls (Fig. 4C). In some fish, we observed a spontaneous development of pigmented tumors in the eye, potentially similar to human ocular melanomas (Fig. 4D). While sample size was small, we observed a reduction in tumor size of more than 30% in all of the ocular melanomas treated with dabrafenib (Fig. 4B, C), suggesting that these tumors are also BRAF-driven, and that dabrafenib treatment has activity towards zebrafish ocular melanoma. Consistent with the rapid absorption and sufficient steady-state levels of tank-water administered dabrafenib (Fig. 1), the treatment resulted in fast and sustained regression of the responding tumors in both skin and ocular melanomas (Fig. 4E). Importantly, tank-water administered dabrafenib was well tolerated with no significant weight-loss (Fig. 4F) or mortality during the two-week treatment. Taken together, tank water -administered dabrafenib is well-tolerated in adult zebrafish over long treatment duration, and results in rapid and sustained tumor regression of BRAF-driven zebrafish melanoma *in vivo*.

**Figure 4.**
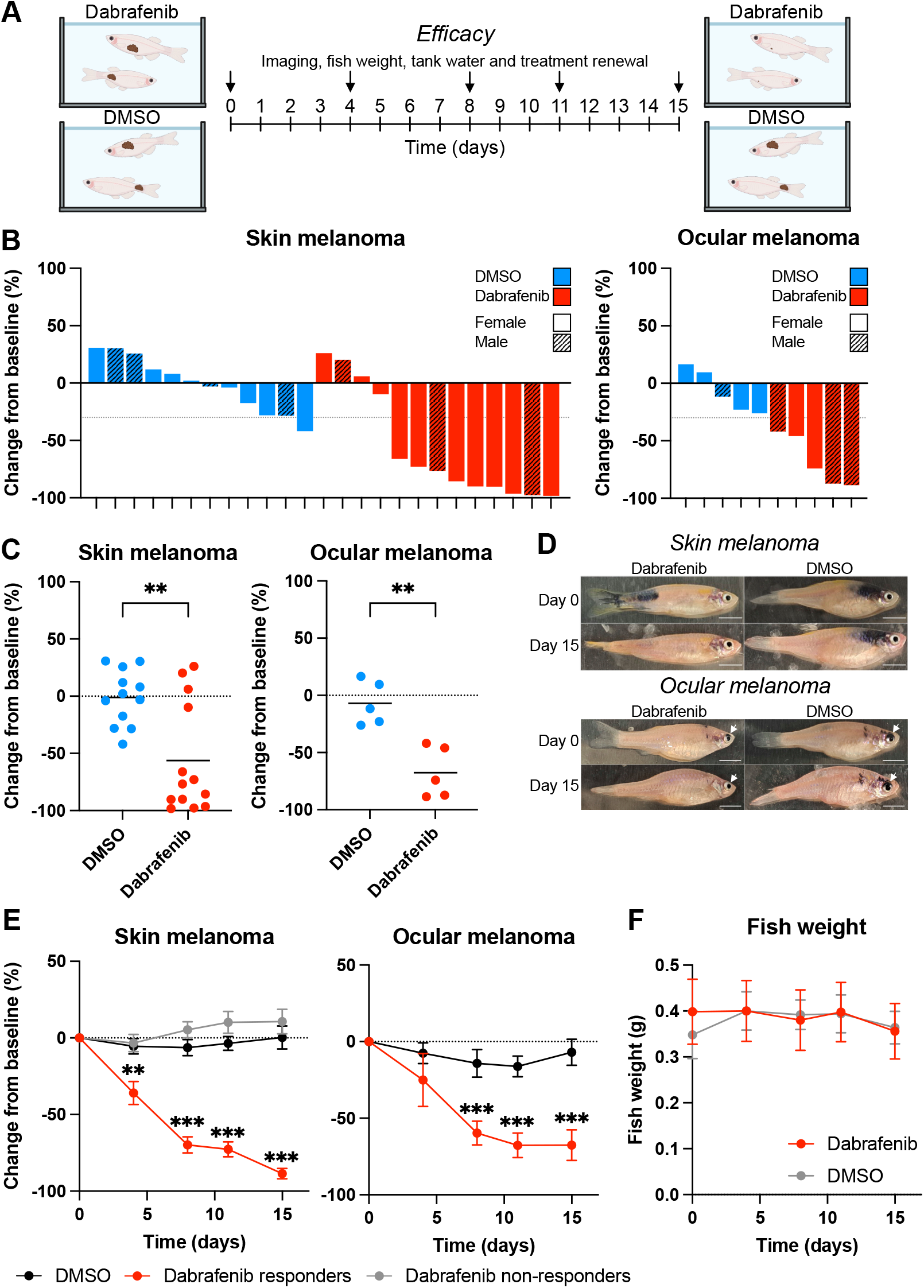
Tank water-administered dabrafenib results in rapid and sustained tumor regression in zebrafish model of BRAF-mutant melanoma. **A)** Schematic presentation of the efficacy experiments. Tumor-bearing fish were treated with 10 µM dabrafenib or DMSO administered in the tank water. The fish were weighed, imaged and the dosing replenished every 3-4 days. The tumor areas of all individual tumors in the study fish were determined from images from all timepoints (see methods). **B)** Waterfall plots demonstrating the tumor area change from baseline after 15 days of treatment for each individual tumor in the study fish. **C)** Scatter plot demonstrating the overall tumor area change from baseline for all tumors in the DMSO or dabrafenib cohorts. **D)** Representative images from DMSO- or dabrafenib-treated fish bearing skin or ocular melanoma. **E)** Tumor area change from baseline over time in DMSO-cohort, as well as tumors in the dabrafenib cohort responding or not responding to treatment. Data shown as mean ± SEM **F)** Fish weight over time in DMSO- or dabrafenib-treated cohorts. Individual unpaired Welch’s t-tests (C), or multiple unpaired Welch’s t-tests with multiple comparisons testing using two-stage step-up method (E) were used to evaluate statistical significance. Asterisks indicate *P*- (C) or *q*- (E) values: ^*^, P < 0.05; ^**^, P < 0.01; ^***^, P < 0.001.

## 3. Discussion

In this work, we studied the suitability of tank administration of BRAF-inhibitor dabrafenib for long-term treatment studies in adult zebrafish. We tested the absorption, distribution, excretion and metabolism of tank-water administered dabrafenib adult fish, as well as the treatment efficacy in genetic zebrafish model of BRAF-mutant melanoma.

Dabrafenib administered to tank water was rapidly absorbed to zebrafish. Plasma levels exceeding in vitro IC50 values were reached within 1 hour of treatment and maintained over the course of the two-week experiment. Dabrafenib also demonstrated fast elimination kinetics, with plasma half-life of 1.6 hours, suggesting rapid metabolism in zebrafish. Indeed, we detected most of the known human metabolites of dabrafenib in zebrafish, suggesting that the metabolism of dabrafenib may follow a similar metabolic process in zebrafish as it does in humans. Consistent with the fast absorption kinetics and therapeutically relevant steady-state plasma levels, tank-water administered dabrafenib led to rapid and sustained tumor regression in a genetic zebrafish model of BRAF-mutant melanoma. We observed responses both in melanomas of the skin, as well as in spontaneously occurring pigmented tumors in the eye, potentially resembling ocular melanomas in humans. In skin melanomas, the treatment resulted similar response rates as observed for human BRAF-mutant melanoma patients treated with BRAF-inhibitors [9,10]. Interestingly, clearly intrinsically resistant tumors were also observed.

Previous reports have described administration of small molecule BRAF inhibitors to zebrafish using gavage [4][5] or feeding [5]. Repeated gavage is a highly technical procedure, requires skilled personnel, and may cause severe morbidity in the treated fish. Both gavage and feeding can be administrated only once or couple of times per day, whereas administration to tank water provides infusion-like continuous administration. A related BRAF inhibitor vemurafenib has been administered in pelleted food once a day with a clear therapeutic response [5]. The pharmacokinetics of vemurafenib were not identified in this study, but at least in humans vemurafenib has a significantly longer half-life [11,12] than dabrafenib [13]. The drug administration *via* food facilitates feasible long-term experiments, but in the case of drugs with short *in vivo* half-life in zebrafish (such as dabrafenib), the administration into tank water may better facilitate the maintenance of effective therapeutic doses throughout long-term treatment. In this regard, repeated administration through injection [14] oral gavage [4] or short daily baths [15] in drug solutions may have similar limitations. The administration directly into tank water is technically easier and well-tolerated method, but the largest drawback is the relatively large amount of drug needed to treat the fish.

The pharmacokinetics is typically understudied in zebrafish and head-to-head comparisons of different delivery routes largely lacking. For future studies, the systematic analysis of pharmacokinetics in this model would be warranted as zebrafish models are increasingly used in drug discovery and development projects, in addition to cancer studies.

To conclude, tank-water administration is a feasible, efficient, and well-tolerated method for treatment of adult zebrafish with dabrafenib, facilitating long-term studies on treatment-induced effects in BRAF-driven tumor biology. Finally, we anticipate that this method also extends to other small molecule inhibitors, especially in the case of molecules with short *in vivo* half-live in zebrafish.

## 4. Materials and Methods

### 4.1. Microinjection of plasmid into zebrafish embryos

Zebrafish of the strain *mitfa(-/-)* and *p53(-/-))*(kind gift of Adam Hurlstone, University of Manchester, UK) were bred and the embryos were collected next morning. Embryos were injected at 1-cell stage with injection volume of 2.3nl of 12.5ng/µl miniCoopR BRAF-V600E-2A-EGFP plasmid, 12.5 ng/µl Tol2TP RNA, 150mM KCl and phenol red in ultrapure water. Injections were done using Nanoject II (Cat:3-000-205A, Drummond) injector and glass needles pulled from glass capillaries (TW100-4, World Precision Instruments Ltd., Sarasota, FL) using micropipette puller (PB-7, Narishige, Tokyo, Japan). After microinjection embryos were maintained in E3 medium (5mM NaCl, 0.17mM KCl, 0.33mM CaCl2, 0.33mM MgSO4) with penicillin-streptomycin at 28.5℃. At 1dpi 20mg/ml pronase E (Cas:9036-06-0, Merck) solution was added to embryos. At 3dpi embryos were selected based on patches or individual dark melanocytes using stereomicroscope.

### 4.2. Zebrafish husbandry

The zebrafish were maintained in accordance to the standard protocols (Westfield, 2000). During the experiments, 5-20-month-old fish were housed in 2 L tanks and were fed with Gemma micro 300 dry food once a day. All experiments were conducted under license obtained from the Animal Experimental Board of the Regional State Administrative Agency for Southern Finland (ESAVI/2688/2020).

### 4.3. Pharmacokinetic analysis from plasma samples

Adult zebrafish (2-3 fish in a 2-litre tank) were treated with 10 µM dabrafenib (HY-14660, MedChemExpress) diluted directly into the tank water. For determining absorption kinetics of dabrafenib, blood samples were taken from fish in different time points after drug administration (0, 1, 2, 4, 8, 24 h and 2 weeks). To determine elimination kinetics, fish were treated for 24 h with 10 µM dabrafenib, followed by transfer to a tank with fresh tank water. Blood samples were taken from fish at different timepoints after transfer to fresh tank water (0, 1, 2, 4, 8, 26 h). Blood samples were collected by transecting the tail between caudal and anal fin with steel blade (Swann-Morton) after chemical euthanasia with 400 mg/l MS222. The tubes and pipette tips were coated with 0.5 M EDTA (pH=8.5, Cas:6381-92-6, VWR Chemicals) in milliQ-water and air-dried. The samples were centrifuged for 5 minutes with a bench-top centrifuge. The plasma was moved to a new EDTA coated tube. Samples were snap-frozen on dry-ice and stored at -80°C.

### 4.4. Quantification of dabrafenib and hydroxy-dabrafenib by LC-MS/MS

#### 4.4.1. Standards and calibration matrix

Blank human plasma was obtained from Blutspendedienst des Bayerischen Roten Kreuzes gGmbH (Munich, Germany). Dabrafenib mesylate (DAB) was purchased from Sigma-Aldrich (Taufkirchen, Germany). Dabrafenib-d9 (DAB-d9; internal standard) was purchased from Alsachim (Illkirch Graffenstaden, France) and Hydroxy-Dabrafenib (HDAB) was purchased from Biozol Diagnostica (Hamburg, Germany).

#### 4.4.2. Stock solutions and working solutions

DAB, HDAB and DAB-d9 stock solutions were prepared in dimethylsulfoxide to yield a concentration of 1 mg/mL. Two independent stock solutions were used for the preparation of calibration standards and quality control (QC) samples. All solutions were stored at -80°C. DAB, HDAB were combined in a working stock solution (solvent: methanol) to yield concentrations of approx. 30 µg/mL for DAB and HDAB. DAB-d9 was diluted in DMSO to 20 µg/mL to get an internal standard (IS) working stock solution. The IS working stock solution was diluted with acetonitrile (ACN) to yield the precipitating agent at an IS concentration of approximately 100 ng/mL DAB-d9. All concentrations refer to the concentration of free base.

#### 4.4.3. Calibration solutions and quality control samples

The calibration curve in matrix was prepared by spiking 950 µL of blank human plasma with 50 µL of the working stock solution to obtain the highest calibration standard. For the highest QC sample, 960 µL of blank human plasma was spiked with 40 µL of the QC working stock solution. The remaining QC samples and calibrators were obtained by serial dilution with blank human plasma. The calibration range for DAB and HDAB was 5-1500 ng/mL. QC high, QC middle and QC low were at a concentration of 1200, 600 and 15 ng/mL, respectively. A weighted calibration (1/conc^2^) was applied for both analytes.

#### 4.4.4. Sample preparation

As the samples contained less than 25 µL of plasma, the available volume was adjusted to 25 µL with blank human plasma. 75 µL of ACN containing IS was added to 25 µL of sample, calibrator or QC sample. After vortexing for 15 s, the samples were centrifuged for 5 min at 4°C and 11.000 rcf. 50 µL of supernatant was transferred to another tube containing 450 µL of a mobile phase mixture (mobile phase A/mobile phase B 60:40, v/v) and vortexed for another 10 s. 300 µL was transferred into a deep-well plate and injected into the HPLC system.

#### 4.4.4. LC-MS/MS experiment

The samples were measured on an API3200 (Sciex, Darmstadt, Germany) combined with an Agilent 1200 HPLC (Agilent, Waldbronn, Germany). The separation was carried out on a Waters XBridge Phenyl column 2.1x50 mm (3.5 µm particle size). The mobile phase A consisted of methanol/water (1:9, v/v) containing 10mM NH_4_HCO_3_ whereas mobile phase B was methanol/water (9:1, v/v) containing 10mM NH_4_HCO_3_. The separation was carried out in gradient separation mode at a flow rate of 400 µL/min. The injection volume was 5 µL. The gradient program was as follows: 40% A / 60% B held for 0.5 min, followed by a linear decrease to 20% A / 80% B over 1.5 min. This composition was maintained isocratically for 3 min, after which the mobile phase was returned to the initial conditions (40% A / 60% B) within 0.25 min and held for 1.75 min to allow column re-equilibration. The total run time was 7 min. Analytes were monitored in positive electrospray ionization (ESI+) mode using multiple reaction monitoring (MRM). For DAB, the transition m/z 520.0 → 307.1 was used with a declustering potential (DP) of 81 V and a collision energy (CE) of 43 V. HDAB was monitored using the transition m/z 536.0 → 323.2 with identical source parameters (DP 81 V, CE 43 V). The internal standard DAB-d9 was monitored using the transition m/z 529.0 → 316.0, also with DP 81 V and CE 43 V. The source parameters where as follows: Turbo IonSpray source (Sciex) at 400 °C and 5500V using source gas 1 and gas 2 at 30 units, curtain gas at 25 units and CAD gas at 6 units.

### 4.5. Mass spectrometry imaging

Zebrafish exposed to with 10 µM dabrafenib for 24 h were snap-frozen in dry ice bath with isopropanol, and moved into –80°C. One fish at a time was embedded on its left side on specimen disk with M-1 Embedding Matrix (Ref:1310, Epredia) inside a cooled Leica CM1950 cryostat, and cut into 15 μm sections. The Mass spectrometry imaging run was made by Turku Metabolomics Centre. 6 dabrafenib-treated samples and 2 control samples from 24-hour timepoint were processed. The matrix was applied using the sublimation method. Super-DHB (Sigma-Aldrich) at a concentration of 40 mg/ml dissolved in acetone was used as a matrix for MALDI. Vacuum was created using the MD 1C +AK+EK chemistry system (VACUUBRAND GMBH). The matrix solution (750 μl) was applied to the bottom of the sublimator chamber and dried under a nitrogen flow, after which a slide with a sample was installed and a vacuum was created. Sublimation was carried out at a temperature of 140°C for 15 minutes.

Samples were measured by Bruker timsTOF fleX equipped with postionization laser (Bruker Daltonics) in positive mode from m/z 300 to 1350 with 100 lasershots per pixel using 50-micron spot size settings (50-micron spot size; 50-micron step size). All imaging experiments were loaded into SCiLS Lab (v2023bPro, Bruker Daltonics) using the default settings with the parameters from FlexImaging. Data were searched for following targets with 10ppm error window: Dabrafenib (DAB, m/z = 520.108), Carboxy-Dabrafenib (CDAB, m/z = 550.083), Desmethyl-Dabrafenib (DDAB, m/z = 506.093), Hydroxy-Dabrafenib (HDAB, m/z = 536.103). The data were not normalized due to scarce distribution of selected targets in analyzed samples. The boxplots were generated within SCiLS Lab (v2023bPro, Bruker Daltonics).

### 4.6. Hematoxylin-eosin staining of mass spectrometry slides

After mass spectrometry imaging, the matrix was removed from samples with 15min incubation in acetone (Cas:67-64-1, Sigma-Aldrich). Samples were incubated twice in 100% ethanol for 3min, once in 70% ethanol for 3 min and once in milliQ-water for 2min. Slides were placed in Mayer’s hematoxylin solution (Ref:MHS16, Sigma-Aldrich) for 10 min followed by 10 min rinse under running tap water, 30 s incubation in milliQ-water and 30 s incubation in 96% ethanol. Then slides were stained with alcoholic Eosin Y solution(Ref:HT110116, Sigma-Aldrich) for 45 s and followed by 1 min wash in 96% ethanol and twice 3min wash in 100% ethanol. The slides were then soaked twice in xylene for 5 min, mounted with DPX mountant (06522, Sigma-Aldrich) and covered with a coverslip. The slides were scanned with Pannoramic P1000 (3DHISTECH) slide scanner with 40x magnification.

### 4.7. Tumour size measurement

The fish of both sexes were housed in groups of 2-3 fish for 16 days. DMSO or 10 µM dabrafenib was administered into the tank water. Three studies at separate times were performed (total n = 28). The fish were visually inspected daily, and nitrogen and oxygen levels were measured every third day to monitor water quality. The fish were weighted and imaged on days 0, 4, 8, 11 and 15 during the experiment under anesthesia with 160 mg/ml tricaine. The tank water and the dabrafenib dosing was renewed at the same time points. The images were taken from both sides and the top of each fish with a mobile phone.

### 4.8. Cell culture

The ZMEL1-GFP cells were cultured in DMEM High glucose media (ECB7501L, Euroclone) containing 10 % fetal bovine serum (FBS) (Biowest), 200 mM glutamine (Euroclone), and 100 U/ml Penicillin /100 ug/ml Streptomycin (Euroclone). The cells were incubated in a humidified incubator at 28°C with 5 % CO2.

### 4.9. Dose-response assay

The ZMEL1-GFP cells were seeded in a 96-well plate at a density of 5,000 cells per well and allowed to adhere for 24 hours. Dabrafenib with concentrations ranging from 1 nM to 10 µM was added to the cells in triplicates using an automatic HD D3000 Digital Dispenser (HP). After 72 h incubation, 30 μl of CellTiter-Glo Luminescent Cell Viability Assay Solution (Promega) was added to each well, and the plate was incubated on shaker (Certomat MO) at room temperature for 10 min in dark. Luminescence was measured using Agilent Biotek Cytation5 Cell imaging multimode reader (AH diagnostics).

### 4.10. Western blotting

The ZMEL1-GFP cells (5x10^6^ cells) were seeded on 10cm dishes. After 24 hours, the cells were treated with dabrafenib concentrations ranging from 1 nM to 10 µM and incubated for further 24 hours. Following treatment, cells were collected in PBS by scraping. After pelleting, the cells were lysed in 150 µl of lysis buffer (150 mM NaCl, 50 mM Tris pH 7.5, 0.4 % Triton-X, 2 mM CaCl2, 2 mM MgCl2, 1 mM EDTA, 5 mM NaF in milliQ-water) supplemented with Halt Protease and Phosphatase Inhibitor Cocktail (ThermoScientific) and Genius Nuclease (Santa Cruz Biotechnology) on ice for 30 min. Protein concentration was determined using the Pierce BCA Protein Assay (Thermo Fisher) according to the manufacturer’s protocol. 30 µg total protein sample was mixed with 6x Laemmli buffer and heated at 95°C for 5 min. Proteins were separated by SDS-PAGE running and transferred onto a nitrocellulose membrane. The membranes were washed with milliQ-water and blocked with 5% milk in TBST (150 mM NaCl, 0.005% Tween-20, 10 mM Tris-HCl pH 7.5 in milliQ-water) for 30 min at room temperature.

Membranes were incubated overnight at 4°C with Phospho-p44/42 MAPK (Thr202/Tyr204) antibody (1:1000, cat# 9101S, Cell Signaling Technology) or p44/42 MAPK antibody (1:1000, cat# 9102S, Cell Signaling Technology) diluted in 5% BSA in TBST with 0.01% NaN3. After washing with TBST, membranes were incubated with IRDye 800CW Goat anti-Rabbit (1:10 000, LI-COR) diluted in blocking buffer for 1.5 h at room temperature. After further TBST washes, signal detection was performed using Odyssey CLx Imager (LI-COR). To control for protein loading, the membranes were stripped with stripping buffer (25 mM glycine-HCl, 1% SDS pH 2) for 20 min and blocked with 5% fat-free milk in TBST before incubating with monoclonal anti-beta-actin antibody (1:1000, cat# A2228, Sigma-Aldrich) and IRDye 680RD Donkey anti-Mouse (1:10 000, LI-COR).

### 4.11. Image analysis

The zebrafish images were analyzed using QuPath (0.3.2). Melanoma lesions from 2-week treatment experiments were quantified by measuring the area of the pigmented tumors. Percentage change for matching pre-, and post-treatment lesions was calculated based on the tumor areas. Response was determined to be over 30% decrease from baseline. The western blot images were quantified using ImageStudio software (5.5) phospho-ERK and ERK band intensities were normalized to actin signal for each lane.

### 4.12. Statistical methods

The statistical analyses were performed and graphs generated using GraphPad Prism 10.4.2. Data is presented as mean ± standard deviation (SD) or standard error of the mean (SEM). For comparisons between two groups, t-test was used. P-values less than 0.05 were considered statistically significant.

## 5. Author contributions

**Jenna Vilman**: Investigation, Methodology, Data curation, Formal analysis, Writing – original draft. **Nikol Dibus**: Investigation, Formal analysis. **Jonas Huhn**: Investigation, Methodology, Writing – review and editing. **Anna-Mari Haapanen-Saaristo**: Investigation, Methodology.

**Ilia Evstafv**: Investigation, Methodology. **Alex Dickens**: Methodology, Resources. **Oliver Scherf-Clavel**: Methodology, Resources, Writing – review and editing. **Ilkka Paatero**: Conceptualization, Methodology, Formal analysis, Funding acquisition, Project administration, Supervision, Writing – original draft, Writing – review and editing. **Kari J. Kurppa**: Conceptualization, Formal analysis, Funding acquisition, Project administration, Supervision, Visualization, Writing – original draft, Writing – review and editing.

### Funding

This work was supported by Research Council of Finland under grant numbers 346656 and 338466 (K.J.K.), Sigrid Jusélius Foundation (K.J.K), Jane and Aatos Erkko Foundation (K.J.K.), Finnish Cultural Foundation (K.J.K.).

## 6. Acknowledgements

We would like to express our gratitude to Zebrafish Core Facility and Metabolomics Core Facility of Turku Bioscience Centre, supported by Biocenter Finland, for their invaluable support, infrastructure and resources. The zebrafish line (*mitfa, p53*) was a kind gift of Dr. Adam Hurlstone (University of Manchester) and ZMEL1-GFP cells were generously granted by Dr. Richard White (Memorial Sloan Kettering Cancer Center, New York, NY, USA).

## References

[1] B.D. Curti, M.B. Faries, Recent advances in the treatment of melanoma., N Engl J Med 384 (2021) 2229–2240. 10.1056/nejmra2034861.

[2] H. Tsao, M.B. Atkins, A.J. Sober, Management of cutaneous melanoma., N Engl J Med 351 (2004) 998–1012. 10.1056/NEJMra041245.

[3] E.E. Patton, H.R. Widlund, J.L. Kutok, K.R. Kopani, J.F. Amatruda, R.D. Murphey, S. Berghmans, E.A. Mayhall, D. Traver, C.D.M. Fletcher, J.C. Aster, S.R. Granter, A.T. Look, C. Lee, D.E. Fisher, L.I. Zon, BRAF Mutations Are Sufficient to Promote Nevi Formation and Cooperate with p53 in the Genesis of Melanoma, Current Biology 15 (2005) 249–254. 10.1016/j.cub.2005.01.031

[4] M. Dang, R.E. Henderson, L.A. Garraway, L.I. Zon, Long-term drug administration in the adult Zebrafish using oral gavage for cancer preclinical studies, DMM Disease Models and Mechanisms 9 (2016) 811–820. 10.1242/dmm.024166.

[5] Y. Lu, E. Elizabeth Patton, Long-term non-invasive drug treatments in adult zebrafish that lead to melanoma drug resistance, DMM Disease Models and Mechanisms 15 (2022) dmm049401. 10.1242/dmm.049401.

[6] OECD, Test Guideline No. 203; Fish, Acute Toxicity Testing, in: OECD Guidelines for the Testing of Chemicals, OECD Publishing, Paris, 2025. http://www.oecd.org.

[7] A. Puszkiel, G. Noé, A. Bellesoeur, N. Kramkimel, M.N. Paludetto, A. Thomas-Schoemann, M. Vidal, F. Goldwasser, E. Chatelut, B. Blanchet, Clinical Pharmacokinetics and Pharmacodynamics of Dabrafenib, Clin Pharmacokinet 58 (2019) 451–467. 10.1007/s40262-018-0703-0.

[8] S. Heilmann, K. Ratnakumar, E.M. Langdon, E.R. Kansler, I.S. Kim, N.R. Campbell, E.B. Perry, A.J. McMahon, C.K. Kaufman, E. Van Rooijen, W. Lee, C.A. Iacobuzio-Donahue, R.O. Hynes, L.I. Zon, J.B. Xavier, R.M. White, A quantitative system for studying metastasis using transparent zebrafish, Cancer Res 75 (2015) 4272–4282. 10.1158/0008-5472.CAN-14-3319.

[9] P.B. Chapman, A. Hauschild, C. Robert, J.B. Haanen, P. Ascierto, J. Larkin, R. Dummer, C. Garbe, A. Testori, M. Maio, D. Hogg, P. Lorigan, C. Lebbe, T. Jouary, D. Schadendorf, A. Ribas, S.J. O’Day, J.A. Sosman, J.M. Kirkwood, A.M.M. Eggermont, B. Dreno, K. Nolop, J. Li, B. Nelson, J. Hou, R.J. Lee, K.T. Flaherty, G.A. McArthur, Improved Survival with Vemurafenib in Melanoma with BRAF V600E Mutation, New England Journal of Medicine 364 (2011) 2507–2516. 10.1056/nejmoa1103782.

[10] A. Hauschild, J.J. Grob, L. V. Demidov, T. Jouary, R. Gutzmer, M. Millward, P. Rutkowski, C.U. Blank, W.H. Miller, E. Kaempgen, S. Martín-Algarra, B. Karaszewska, C. Mauch, V. Chiarion-Sileni, A.M. Martin, S. Swann, P. Haney, B. Mirakhur, M.E. Guckert, V. Goodman, P.B. Chapman, Dabrafenib in BRAF-mutated metastatic melanoma: A multicentre, open-label, phase 3 randomised controlled trial, The Lancet 380 (2012) 358–365. 10.1016/S0140-6736(12)60868-X.

[11] J.F. Grippo, W. Zhang, D. Heinzmann, K.H. Yang, J. Wong, A.K. Joe, P. Munster, N. Sarapa, A. Daud, A phase I, randomized, open-label study of the multiple-dose pharmacokinetics of vemurafenib in patients with BRAF V600E mutation-positive metastatic melanoma, Cancer Chemother Pharmacol 73 (2014) 103–111. 10.1007/s00280-013-2324-5.

[12] K.T. Flaherty, I. Puzanov, K.B. Kim, A. Ribas, G.A. Mcarthur, J.A. Sosman, P.J. O’dwyer, R.J. Lee, J.F. Grippo, K. Nolop, P.B. Chapman, M. General, Inhibition of Mutated, Activated BRAF in Metastatic Melanoma, N Engl j Med 363 (2010) 809–828. 10.1056/NEJMoa1002011

[13] G.S. Falchook, G. V Long, R. Kurzrock, K.B. Kim, T.H. Arkenau, M.P. Brown, O. Hamid, J.R. Infante, M. Millward, A.C. Pavlick, S.J. O’day, S.C. Blackman, C.M. Curtis, P. Lebowitz, B. Ma, D. Ouellet, R.F. Kefford, Dabrafenib in patients with melanoma, untreated brain metastases, and other solid tumours: a phase 1 dose-escalation trial, Lancet 379 (2012) 1893– 901. 10.1016/S0140-6736(12)60398-5

[14] M.D. Kinkel, S.C. Eames, L.H. Philipson, V.E. Prince, Intraperitoneal injection into adult zebrafish, Journal of Visualized Experiments (2010) e2126. 10.3791/2126.

[15] A.T. Monstad-Rios, C.J. Watson, R.Y. Kwon, ScreenCube: A 3D Printed System for Rapid and Cost-Effective Chemical Screening in Adult Zebrafish, Zebrafish 15 (2018) 1–8. 10.1089/zeb.2017.1488.

